# Structural transitions in Orb2 prion-like domain relevant for functional aggregation in memory consolidation

**DOI:** 10.1101/2020.07.08.193656

**Authors:** Javier Oroz, Sara S. Félix, Eurico J. Cabrita, Douglas V. Laurents

## Abstract

The recent structural elucidation of *ex vivo Drosophila* Orb2 fibrils revealed a novel amyloid formed by interdigitated Gln and His residue side chains belonging to the prion-like domain. However, atomic-level details on the conformational transitions associated with memory consolidation remain unknown. Here, we have characterized the nascent conformation and dynamics of the prion-like domain (PLD) of Orb2A using a nonconventional liquid-state NMR spectroscopy strategy based on ^13^C detection to afford an essentially complete set of ^13^Cα, ^13^Cβ, ^1^Hα and backbone ^13^CO and ^15^N assignments. At pH 4, where His residues are protonated, the PLD is disordered and flexible, except for a partially populated α-helix spanning residues 55-60. At pH 7, in contrast, His residues are predominantly neutral and the Q/H segments adopt minor populations of helical structure, show decreased mobility and start to self-associate. At pH 7, the His residues also bind Zn^++^, which promotes further association. These findings represent a remarkable case of structural plasticity, based on which an updated model for Orb2A functional amyloidogenesis is advanced.

**Highlights:**

· The Orb2 prion like domain that forms the structures related to memory consolidation is studied by solution NMR.
· The amyloidogenic Q/H-rich stretch is disordered and flexible at low pH.
· Residues 55-60 form a partly populated α-helix at pH 4.
· At pH 7, the Q/H-rich segment also adopts a low population of α-helix and rigidifies.
· Zn^++^ binding induces associative changes in the Orb2 prion-like domain.

## Introduction

The comprehension of how the nervous system encodes memory has been an important goal since the beginning of modern neuroscience. At the turn of the twentieth century, memory was proposed to be encoded as alterations in the neurite network (Cajal, 1894) as a persistent chemical change (Semon, 1904). This hypothesis has gained support over the decades (Hebb, 1949; Frey & Morris, 1997; Kandel, 2006; Josselyn & Tonegawa, 2020), and in 2003, Si, Lindquist and Kandel published a seminal paper providing the first evidence for the molecular basis of memory consolidation (Si *et al*., 2003). In that study, Si and coworkers proposed that functional aggregation of the protein CPEB into ordered structures is key to memory consolidation in *Aplysia*. Numerous studies have corroborated the key functional role of CPEB aggregation as well as that of its homologs in memory consolidation in *Drosophila* and mammals (Si & Kandel, 2016).

The *Drosophila* CPEB homolog, named Orb2, has been investigated in the most detail. Two Orb2 isoforms, called Orb2A and Orb2B, are relevant for long term memory in *Drosophila melanogaster* (Keleman *et al*., 2007; Krüttner *et al*., 2012). Whereas they differ at the N-terminus, both Orb2A and Orb2B contain a prion-like domain (PLD) rich in Gln and His residues, followed by a stretch of Gly and Ser residues (**Sup. Fig. 1A**). Whereas the Orb2A isoform is rare, it is essential for triggering the aggregation of the more abundant Orb2B and to maintain the memory trace (Krüttner *et al*., 2012; Majumdar *et al*., 2012). The first nine residues of Orb2A are unique to this isoform (**Sup. Fig. 1A**). Rich in hydrophobic side chains, these nine residues have been reported to interact with membrane mimetics (Soria *et al*., 2017), to be crucial for aggregation (Majumdar *et al*., 2012) and to adopt a β-strand within an Orb2 amyloid structure formed *in vitro* as characterized by solid-state NMR and EPR (Cervantes *et al*. 2016).

Both Orb2A and Orb2B contain a 31-residue long Q/H-rich stretch, followed by a short amphiphilic segment (residues N_55_LSAL_59_) and a second modest Q/H-stretch (H_60_HHHQQQQQ_68_) (**Sup. Fig. 1A**). Due to its similarity to amyloid-forming polyQ stretches in huntingtin (Scherzinger *et al*., 1997) and the androgen receptor (Eftekharzadeh *et al*., 2016), the main Q/H-rich stretch (residues 23-53) was suspected to be key for Orb2 amyloid formation. Indeed, while both Q/H-rich stretches appear rather disordered in Orb2 amyloids formed *in vitro* (Cervantes *et al*. 2016), Orb2 aggregation and memory consolidation can be blocked in *Drosophila* by a peptide which inhibits polyglutamine aggregation (Hervás *et al*., 2016). Very recently, the elucidation of the structure of physiological Orb2 amyloid isolated directly from *Drosophila* fly brains has shown that the main Q/H-rich stretch does in fact form the physiologically relevant amyloid structure (Hervás *et al*., 2020a). This amyloid features three-fold symmetry and side chain to mainchain H-bonds (**Sup. Fig. 1B**). The participation of numerous His residues in the amyloid core provides a singular mechanism for amyloid destabilization through acidification. Intriguingly, the first nine residues of Orb2A, while relevant for triggering aggregation (Majumdar *et al*., 2012), do not contribute to the amyloid core, which is composed of the more abundant Orb2B isoform (Hervás *et al*., 2020a) (**Sup. Fig. 1**).

Following the second Q/H-rich stretch, there is a two-hundred residue, presumptively disordered region with an elevated content of Gly, Ser, Pro and Asn (**Sup. Fig. 1A, Sup. Fig. 2**). The conserved C-terminal half of Orb2 is composed of two RRM domains, which bind Orb2-specfic mRNA targets, and a ZZ-type zinc finger domain (**Sup. Fig. 1A**). ZZ-type zinc fingers are a special sub-class that bind two Zn^++^ ions and generally mediate protein/protein interactions, not nucleic acid binding (Legge *et al*., 2004). In Orb2, the conserved C-terminal domains are crucial for binding mRNAs containing a uridine-rich 3’UTR (Mastushita-Sakai *et al*., 2010) and potentially different translation complexes (Hervás *et al*., 2020a). This binding suppresses the translation of mRNAs coding for factors promoting synapse formation, synapse growth and proteases (Mastushita-Sakai *et al*., 2010). Upon neuronal stimulation, Orb2 forms amyloid leading to the activation of these mRNAs (Khan *et al*., 2015). Due to its physiological importance, Orb2 aggregation is finely controlled. Studies have reported key roles of intron retention (Gill *et al*., 2017), phosphorylation and protein stability (White-Grindley *et al*., 2014) and chaperones (Li *et al*., 2016) in tightly regulating Orb2 functional amyloid formation. Besides the mentioned control of Orb2 aggregation via acidification (Hervás *et al*., 2020a), Siemer and co-workers proposed that the spacing of His residues within the long Q/H-rich stretch would position them on the same face of a hypothetical α-helix (Bajakian, *et al*., 2017). This may contribute to the binding of Ni^++^, Cu^++^ and Zn^++^ which might impact aggregation (Bajakian, *et al*., 2017). Interestingly, the longer Q/H-rich region of Orb2 may be sequestered into pathological amyloids formed by expanded polyQ segments of huntingtin, providing a novel explanation for memory loss in dementia (Hervás *et al*., 2016; Joag *et al*., 2019).

Whereas all these results have provided molecular insight into Orb2’s role in memory consolidation, much is still unclear regarding the atomic level details of the amyloid formation process. In particular, it is unknown whether the nascent protein contains elements of partial structure which predispose the formation of the functional amyloid, or alternatively, act as a safety mechanism to prevent premature or excess amyloid formation. The objective of this study is to characterize the atomic level conformation and dynamics of Orb2A PLD and the first residues of the G/S-rich region using solution NMR spectroscopy. Because pH is proposed to impact amyloid formation and stability (Hervás *et al*., 2020a), we have characterized the conformation of Orb2A both at pH 4, where His residues are positively charged which impedes aggregation, as well as pH 7, where neutral His are compatible with amyloid formation. In addition, the binding of Zn^++^ promotes self-association via the Q/H-rich regions. The structural transitions observed may represent nascent structures in the process of aggregation into functional amyloids.

## Results

### The Orb2A PLD is disordered at pH 4.0, except for a partly populated α-helix spanning residues 55-60

Initial 1D ^1^H and 2D ^1^H-^15^N HSQC spectra recorded in PBS at pH 7.0 revealed broad lines which are not optimal for spectral analysis and assignment. Therefore, following the low pH, low ionic strength strategy of Song and coworkers (Li *et al*., 2006), which we have recently applied to assign the snow flea antifreeze protein (Treviño *et al*., 2018) and a RepA winged helix domain (Pantoja-Uceda *et al*., 2020), spectra of Orb2A PLD were recorded at pH 4.0 in 1 mM acetic acid buffer at 25 °C. Under these conditions, the signals were sharper and gave excellent quality 2D and 3D spectra. The ^1^Hα, ^1^HN, ^13^Cα, ^13^Cβ and backbone ^15^N and ^13^CO chemical shifts recorded at pH 4.0, 25 °C have been deposited in the BMRB data bank under accession number **50274**. The assignments are complete except for the ^1^Hα of proline residues and the ^13^CO and ^13^Cβ of terminal Ser 88.

The assigned 2D ^1^H-^15^N HSQC and 2D ^13^C-^15^N CON spectra registered at pH 4.0, 25 °C are shown in **Figure 1A** and **1B**, respectively. The resonances are generally sharp and well defined except for clusters of overlapped Gln and His residues belonging to the amyloidogenic tract (**Sup. Fig. 1B**). The low ^1^HN signal dispersion is a hallmark of disordered proteins and an assessment of the ^13^Cα, ^13^Cβ, ^13^CO and ^15^N chemical shifts using the TALOS+ program confirmed the overall dearth of structure in the Orb2A PLD under these conditions. With regard to the glutamine side chain amide groups, their ^1^H_2_-^15^N signals are also clumped into overlapped peaks (**Sup. Fig. 3**). Notably, they do not show the striking pattern of disperse signals observed for the glutamine side chain resonances of the androgen receptor, which participate in side chain to backbone hydrogen bonds (Escobedo *et al*., 2019). Orb2A PLD contains a single Cys residue. Its ^13^Cβ chemical shift indicates that it is reduced at pH 4.0 in 1 mM acetic acid buffer in the absence of reducing agent. Regarding the proline residues, all Xaa-Pro peptide bonds are in the *trans* conformation, which is expected for a disordered chain, as judged by their ^13^Cβ and ^13^CO chemical shifts.

**Figure 1:**
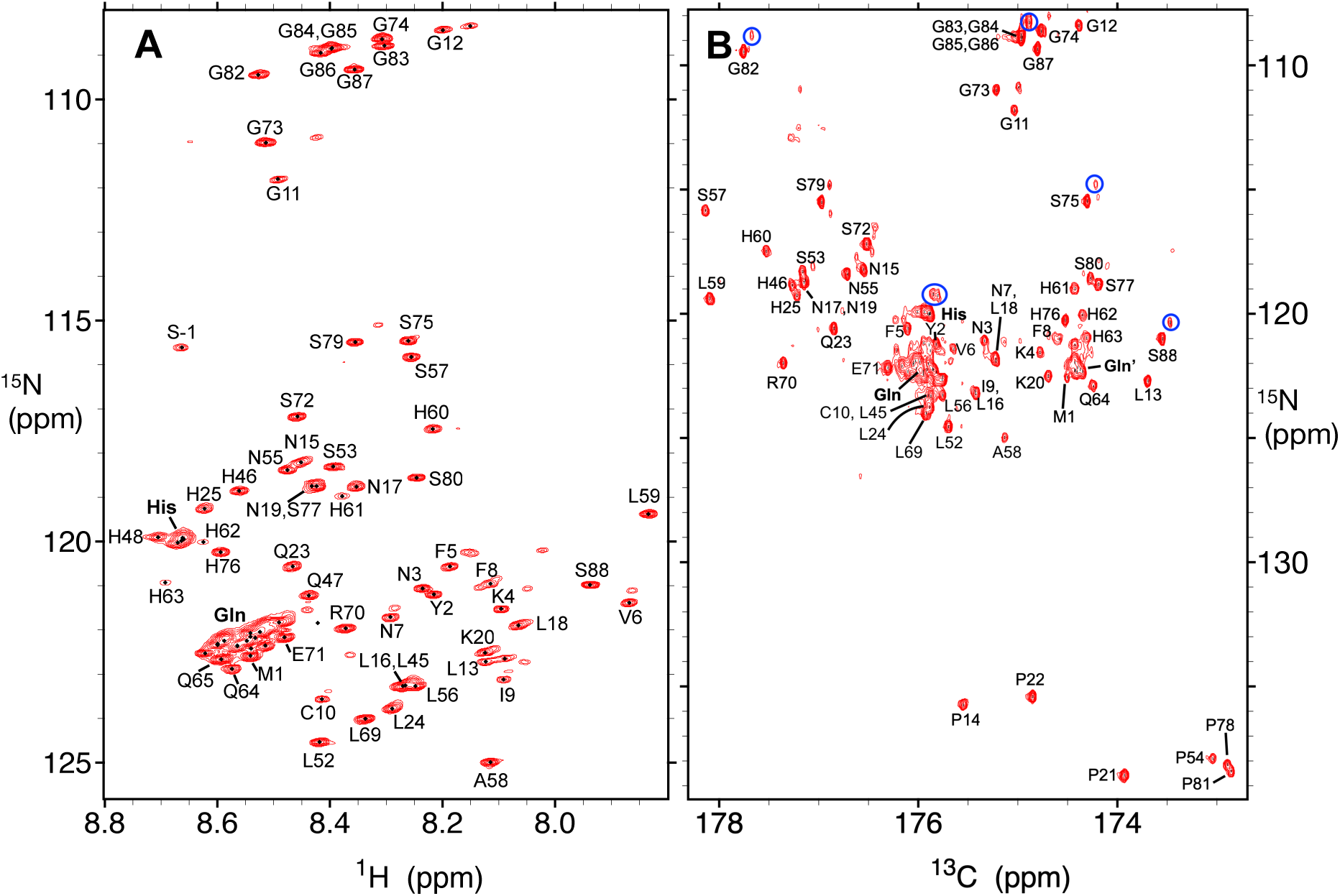
Assigned 2D ^1^H-^15^N HSQC and 2D ^13^C-^15^N CON spectra of Orb2A PLD recorded at pH 4.0, 25°C. **A.** 2D ^1^H-^15^N HSQC spectrum of Orb2A PLD recorded at pH 4.0, 25°C. Individual assigned signals are labeled. S-1 corresponds to the Ser residue resulting from cloning and tag cleavage that precedes M1. The bold **His** and **Gln** labels refer to overlapped His and Gln peaks; these arise from residues in the amyloidogenic segment. The multiplication factor between contour levels is 1.4 here and in panel **B** as well as **Figure 4A**. **B.** 2D ^13^C-^15^N CON spectrum of Orb2A PLD recorded at pH 4.0, 25°C. The labels shown correspond to the ^15^N signals; the ^13^CO correlations arise from the preceding residue. Note that the y-axis scale is longer in panel **B** than **A** to include Pro residues. The bold **His, Gln** and **Gln’** labels are for overlapped His and Gln residues; the **Gln’** cluster comes from *i* His ^13^CO /*i+1* Gln ^15^N correlations. For intense or overlapped peaks, a satellite peak is often seen slightly to the right and above the main peak. These satellite peaks arise from deuteration and some examples are indicated with **blue** circles.

The ^13^Cα, ^13^Cβ and ^13^CO Δδ values, which reveal partially populated elements of secondary structure, are plotted in **Figure 2A**. The segment composed of residues 55-60, which follows the amyloidogenic Q/H-rich stretch, adopts a minor population of α-helix. Based on the conformational chemical shift (Δδ values expected for 100% α-helix of 3.1, - 0.4 and 2.2 ppm for ^13^Cα, ^13^Cβ and ^13^CO, respectively (Spera & Bax, 1991; Wishart & Skyes 1994), the population of α-helix for these residues is approximately 20%. Otherwise, the N-terminal hydrophobic stretch, the Q/H-rich segments and the C-terminal G-rich element do not contain detectable populations of secondary structure under these conditions. Whereas the Δδ 13CO values of terminal G_82_GGGGG_87_ are anomalously high at pH 4 (and also pH 7, see below), which suggests α-helix formation, they are not corroborated by high Δδ ^13^Cα values, so this may rather reflect a shortcoming in the prediction of secondary structure from ^13^CO δ values for polyG repeats. As an orthogonal measure of secondary structure, the 3J_HNHA_ coupling constants were measured and analyzed using the Karplus equation to obtain backbone ϕ angles. Some ^3^J_HNHA_ coupling constants appear to be somewhat lower for residues 55-60 (**Figure 2B**) which may reflect the minor population of α-helix. Next, the μs/ms dynamics were assessed by R_1_ρ relaxation measurements. The obtained values are consistent with high mobility, except for residues 55-60, whose slightly elevated R_1_ρ rates reflect a modestly increased rigidity, due to the partial α-helical structure (**Figure 2C**).

**Figure 2:**
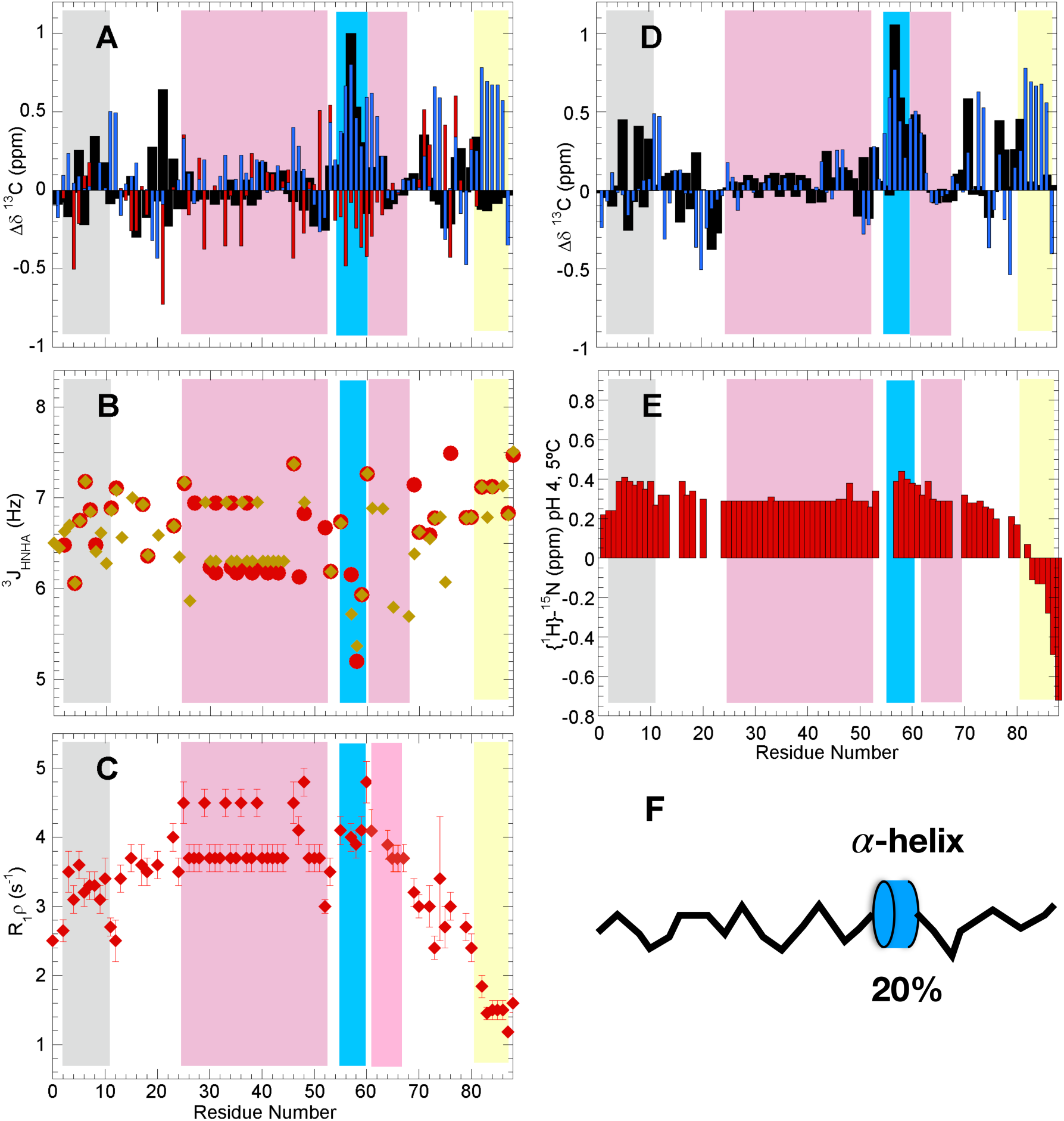
Orb2A PLD is largely disordered pH 4, save a short α-helix in residues 55-60. In all panels, the different zones of the Orb2 PLD are shaded as follows: N-terminal hydrophobic stretch = **gray**, Q/H-rich = light magenta, residues 55-60= **blue**, H·Q-rich= light magenta, G-rich=**yellow**. In the Q/H or H·Q-rich regions most of the values shown come from averaging of overlapped His or Gln peaks. **A.** Conformational chemical shifts (Δδ) of Orb2 PLD at pH 4, 25 °C. ^13^Cα= **black** bars, ^13^CO= **blue** bars, ^13^Cβ= **red** bars. Experimental uncertainties are ≤ ± 0.04, 0.02 and 0.08 ppm for ^13^Cα, ^13^CO & ^13^Cβ, respectively. **B.** ^3^J_HNHA_ coupling constants at pH 4, 25°C. Values of 5 Hz or less are indicative of α-helix; higher values coil or extended conformations. Values shown in **gold** were measured by peak height using the Sparky program. Values in **red** were obtained using the peak integral function of Topspin 4.0. **C.** Transverse relaxation rates in the rotating frame (R_1_ρ) at pH 4.0, 25 °C. **D.** Conformational chemical shifts (Δδ) of Orb2 PLD at pH 4, 5 °C. ^13^Cα= **black** bars, ^13^CO= **blue** bars. The experimental uncertainties are generally ≤ ± 0.05 ppm for both ^13^Cα and ^13^CO. **E.** {^1^H}-^15^N NOE at pH 4.0, 5.0°C. Values approaching 0.85 indicate stiffness on fast ps/ns and are typical of well-folded rigid proteins; values less than 0 indicate a high flexibility. Uncertainties are about ± 0.01. **F.** Schematic diagram of Orb2 PLD at pH 4 featuring a flexible and disordered conformational ensemble with a short, modestly populated α-helix spanning residues 55 – 60.

The population of marginally stable elements of secondary structure in polypeptides is generally enhanced by cooling. To check if this is the case for Orb2A PLD, additional spectra were recorded at 15°C and 5°C, pH 4.0. Nevertheless, the analysis of the conformational chemical shifts did not reveal any additional elements of secondary structure, beyond the 55-60 helix already detected at 25°C (**Figure 2D**). Finally, the fast ps/ns dynamics were characterized by {^1^H}-^15^N NOE measurements at 5°C. Low values indicative of flexibility are seen throughout the domain and are only slightly higher for the hydrophobic segment and α-helix spanning residues 55-60 (**Figure 2E**). The G/S-rich C-terminal residues are especially mobile. Taking all these results together, we can conclude that at pH 4.0, where the His residues of Orb2A’s PLD are chiefly positively charged, the polypeptide chain is disordered and flexible, except for the minor population of α-helix formed by residues 55-60 (**Figure 2F**).

### The amyloidogenic Q/H-rich segment adopts partly populated helical conformations and rigidifies at neutral pH

A visible precipitate formed when the pH of the sample was increased from 4.0 to 7.0. Nevertheless, the concentration of protein remaining in solution was sufficient to record good quality 2D ^1^H-^15^N and ^1^H-^13^C HSQC as well as 3D ^1^H-detected HNCO, HNCA and CBCAcoNH spectra. The 2D ^1^H-^15^N HSQC spectrum (**Figure 3A**) is significantly broadened. The average ^1^HN peak width is 27 ± 4 Hz at pH 7.0 compared to 17 ± 2 Hz at pH 4.0 (**Figure 3B**). This is an indication of conformational interconversion, oligomerization, or both (Charlier *et al*., 2016). The assignments at pH 7.0 are complete for ^13^Cα, ^13^Cβ, ^1^HN and backbone ^13^CO and ^15^N nuclei save the ^13^Cβ of S77, the ^13^CO of S79, the ^13^Cβ and ^13^CO of S88 as well as the ^13^Cβ and ^13^CO of L15 and other residues preceding proline residues. Moreover, all assignments are missing for H63, H64 and H78 as these residues’ ^1^HN signals may be adversely affected by exchange broadening. The assigned chemical shift values, like those measured at pH 4.0, have been deposited in the BMRB under assession number **50274**.

**Figure 3:**
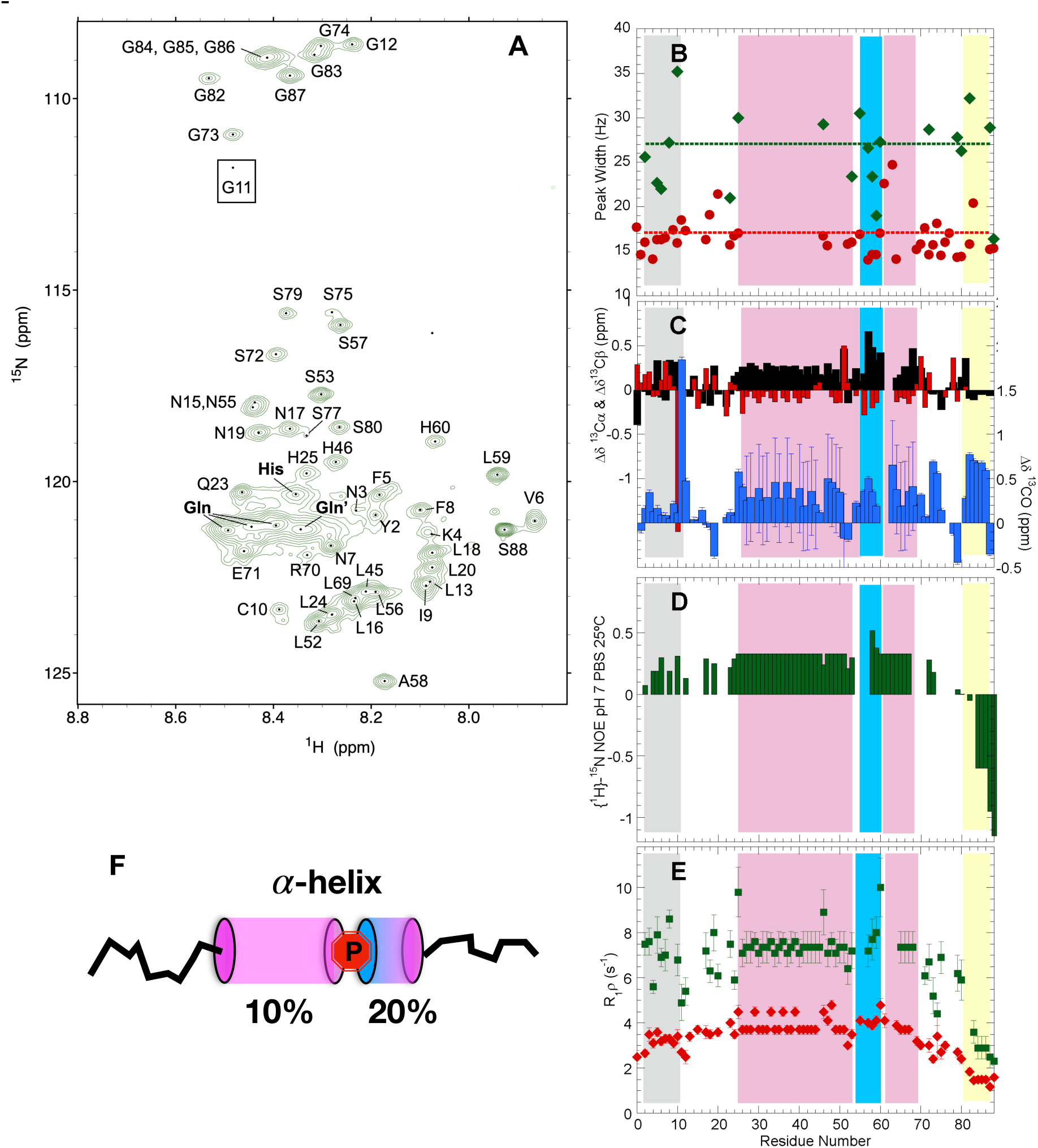
The Q/H-stretch of Orb2A partially adopts α-helical conformations and rigidifies at pH 7. In panels **B-E**, the different zones of the Orb2 PLD are shaded as follows: N-terminal hydrophobic stretch = **gray**, Q/H-rich = **light magenta**, residues 55-60= **blue**, H·Q-rich= **light magenta**, G-rich=**yellow**. In the Q/H or H·Q-rich regions most of the values shown come from averaging of overlapped His or Gln peaks. **A.** 2D ^1^H-^15^N HSQC spectrum of Orb2A PLD registered at pH 7.0, 25°C. The bold **His, Gln and Gln’** labels refer to overlapped His and Gln peaks; these arise from residues in the amyloidogenic segment. The Gln’ peak arises from Gln which follow a His along the sequence. The signal of G11 (*boxed*) is beneath the lowest selected contour level. To afford a fair appreciation of the wider peak widths observed here relative to the spectrum at pH 4, the same axes, aspect ratio and multiplication factor between contour levels (1.4) are used here as in **Figure 2A**. **B**. ^1^H peak width of Orb2 PLD ^1^H-^15^N HSQC peaks at 25 °C and pH 7 (**green** diamonds) and pH 4 (**red** circles). The dotted lines represent the average peak widths at pH 7 (**green**) and pH 4 (**red**). Overlapped peaks are excluded. The uncertainties are approximately ± 1.5 Hz. **C.** Conformational chemical shifts for ^13^Cα (**black** bars) ^13^Cβ (**red** bars) and ^13^CO (**blue** bars) at pH 7.0, 25°C. For clarity, the y-axis scale is shifted for the ^13^CO Δδ values. The ^13^CO of Gly 11 and the ^13^Cβ of Cys are outliers. The ^13^CO Δδ values of the Q-rich region have large uncertainties due to the broad nature of the overlapped peak of Q residues. **D.** Fast ps/ns dynamics assessed by the heteronuclear {^1^H}-^15^N NOE. Uncertainties are about ± 0.015. **E.** Transverse relaxation rates in the rotating frame (R_1_ρ) for Orb2 PLD at pH 7.0, 25°C (**green** squares). For comparison, the R_1_ρ value obtained at pH 4.0, 25 °C and already shown in **Fig. 3C** are also shown here by open **red** diamonds. **F.** At pH 7, the Q/H segments (**magenta** cylinders) and the 55-60 segment (**blue** cylinder) adopts a small but detectable population of α-helical conformers separated by proline 54 (**red**).

Remarkably, the assessment of the Δδ values shows that at neutral pH, the main Q/H-rich stretch partly adopts α-helical conformations, whose population is approximately 10% (**Figure 3C**). Residues S_53_-P_54_ separate this helix from that formed by residues 55-60. The shorter Q/H segment also appears to adopt a minor population of α-helix. The nine N-terminal residues do not show a clear tendency to adopt a preferred secondary structure at pH 7. Cys10 remains reduced and Xaa-Pro peptide bonds continue to be mainly *trans* at pH 7. Exchange measurements at pH 7 yielded {^1^H}-^15^N NOE and R_1_ρ values which are elevated for most residues (**Figure 3D, 3E**) relative to values obtained at pH 4.0. These measurements reflect stiffening on ps/ns and μs/ms timescales and is consistent with the formation of preferred conformers and associative processes. However, the G/S-rich segment located C-terminal to the PLD continues to be highly dynamic at pH 7, both on faster ps/ns timescales as well as slower μs/ms timescales.

After a two month incubation at 4 °C, Orb2A PLD’s ^1^H-^15^N NMR spectra at 25°C in PBS revealed that despite some changes, most peaks retained their positions; this suggest that slow conformational changes or aggregative processes are minimal (**Fig. 4A**). Zinc chloride was then added to a final concentration of 2 mM. In the presence of this divalent cation, the peak intensities of His and Gln ^1^H-^15^N resonances drop practically to the signal/noise limit (**Fig. 4B, C**). Strong decreases are also seen for the ^1^H-^13^Cβ and ^1^H-^13^C Cα signals of His in 2D ^1^H-^13^C HSQC spectra (**Sup. Fig. 4**). This strongly suggest that Zn^++^ binds to His residues and promotes association processes of nearby Gln residues which are manifested as a loss of signal intensity. The signal intensity of other residues also decreased following Zn^++^ addition, but sufficed to record good quality 3D HNCO, HNCA and CBCAcoNH spectra. The analysis of the these spectra revealed that the conformational ensemble of the molecules remaining visible to NMR is poor in helical structures and slightly enriched in extended conformations (**Fig. 4D**). Thus, Zn^++^ binds the His residues and drives the association of helical conformations of Orb2A PLD into large assemblies invisible to liquid state NMR.

**Figure 4:**
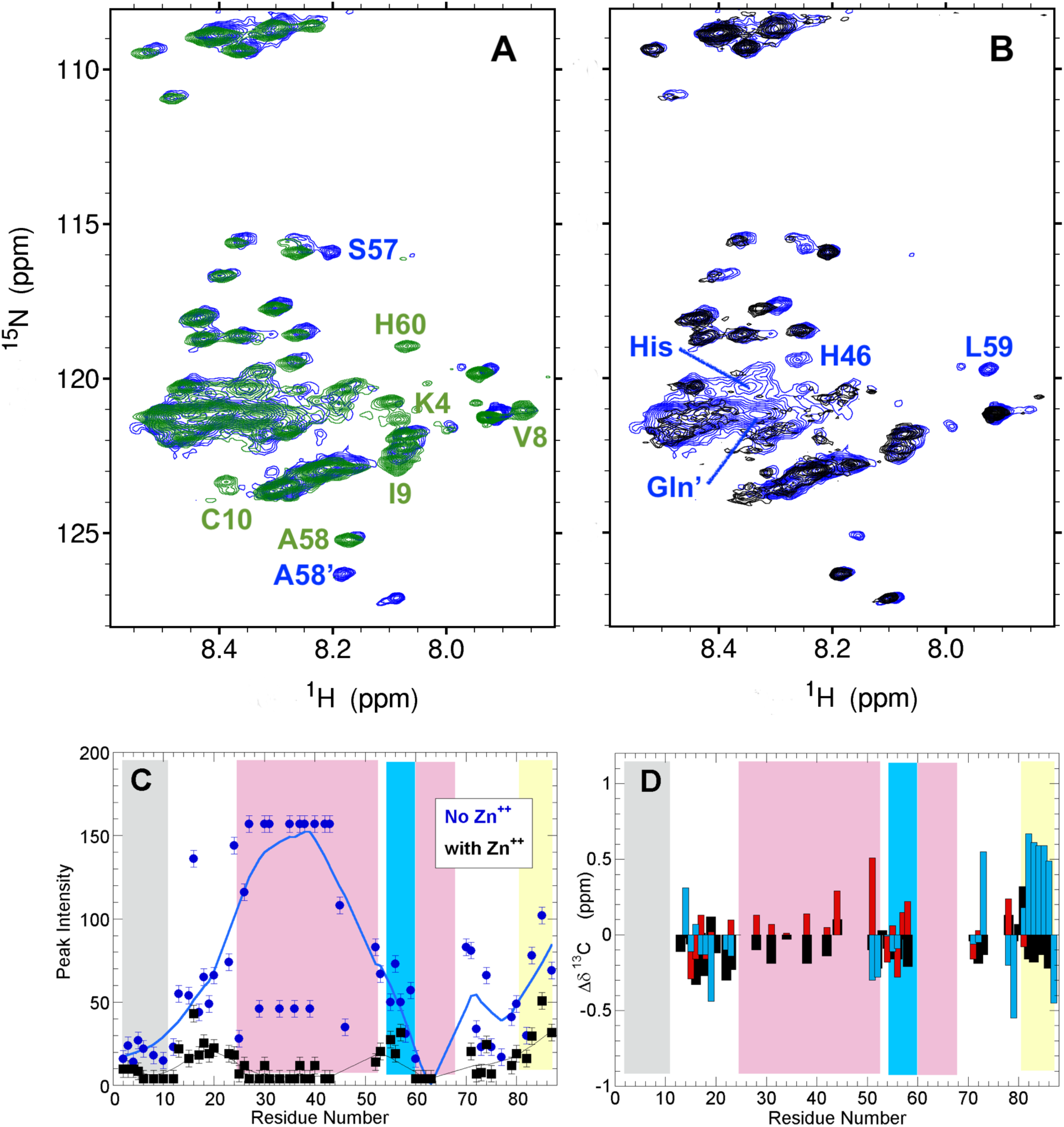
Zn^++^ binding to His residues promotes further oligomerization. **A.** ^1^H-^15^N HSQC before (**green**) and after (**blue**) the 2 month pause. Some distinct residues are labeled. **B.** ^1^H-^15^N before (**blue**) and after **(black**) the addition of ZnCl_2._. His and Gln signals (labeled) weaken sharply after addition of Zn^++^. **C.** The largest drop in peak intensity upon Zn addition occurs for residues of the hydrophobic segment (**gray** band) and the Q/H-rich segments (**rose** band). **D.** Conformational chemical shifts for ^13^Cα (**black**), ^13^Cβ (**red**) and ^13^CO (**blue**) strongly suggest that the molecules remaining in solution lack helical zones and are disordered.

## Discussion

The key findings of this study are that the amyloidogenic Q/H-rich region of Orb2’s PLD is disordered and flexible at low pH, but adopts a minor population of α-helix at pH 7.0, where the His residues are mostly in the neutral state. The hypothesis that Orb2 aggregation triggers enduring synapse-specific alterations in translation which are key for memory consolidation received strong support by the elucidation of its amyloid structure from adult *Drosophila* heads (Hervás *et al*., 2020a). Gln and His side chains are interlocked in this amyloid (**Sup. Fig. 1B**). The relative instability of Orb2 amyloid at acidic conditions, as shown in Sup. Fig. 15 of Hervás *et al*., 2020a, can be attributed to His side chains becoming positively charged and producing electrostatic repulsion at low pH. *In vivo*, this may facilitate the disintegration of Orb2 amyloid fibrils in the lysosome. Considering the results reported here, is it possible that Orb2 PLD histidines could be initially maintained in a charged state *in vivo*, and then neutralized to help trigger amyloid formation?

In the *Drosophila* synapse, Orb2 monomers are initially bound to target mRNA molecules via their RRM domains and the proximity of polyanionic mRNA, phosphorylated Tob (White-Grindley *et al*., 2014), and/or possibly other anions of the neuronal granule (Ford *et al*., 2019) could tend to stabilize the charged imidazolium form of His side chains. Upon neural stimulation, the influx of Ca^++^ and Zn^++^ cations, which are excellent imidazole ligands, could displace the imidazolium H^+^ and bind neutral His. This scenario is supported by ITC measurements demonstrating that the Q/H-rich region of Orb2 binds divalent metal cations (Bajakian *et al*., 2017), which we corroborate here by NMR for Zn^++^ (**Figure 4**). Those researchers also proposed that if this segment were to adopt an α-helical conformation (just as we have shown here), then the His residues, which are mostly spaced *i, i+3* and *i, i+4*, would be mainly positioned on the same side of the α-helix (Bajakian *et al*., 2017). The binding of cations such as Zn^++^ to such an array of His residues could increase the formation of α-helix, although this point is difficult to test due to the Orb2A PLD aggregation that accompanies cation binding, even at low concentrations used for CD spectroscopy (Bajakian *et al*., 2017). The resulting helical conformations may drive association processes such as the formation of coiled-coils, as seen for other proteins which undergo liquid-liquid phase separation via intermolecular helical associations (Conicella *et al*., 2016). Q-rich coiled-coils and higher oligomeric species have been proposed as intermediates in amyloidogenesis of *Aplysia* and mammalian CPEBs (Fiumara *et al*., 2010; Pelassa *et al*., 2014; Hervás *et al*, 2020b; Ramírez de Mingo, *et al*., 2020a), which are probably heterogeneous as strongly suggested by kinetic modeling (Vitalis and Pappu, 2011). A hypothetical working model which incorporates these ideas is shown in **Figure 5**. Whereas *Aplysia* and mammalian CPEB homologs share a similar domain organization with *Drosophila* Orb2, they lack the multiple His residues interspersed in their Q-rich segments (Hervás *et al*, 2020b; Ramírez de Mingo, *et al*., 2020a) which are found in Orb2. This suggests that the pH-regulated nature of Orb2 amyloid formation and dissociation may be substituted by other mechanisms in higher organisms. Proline 54, which interrupts the helices formed by the Q/H-rich segments and residues 55-60, may play an important role in preventing premature amyloid formation. Similar proline residues which delimit α-helices have been recently described in human CPEB3 (Ramírez de Mingo *et al*., 2020a).

**Figure 5:**
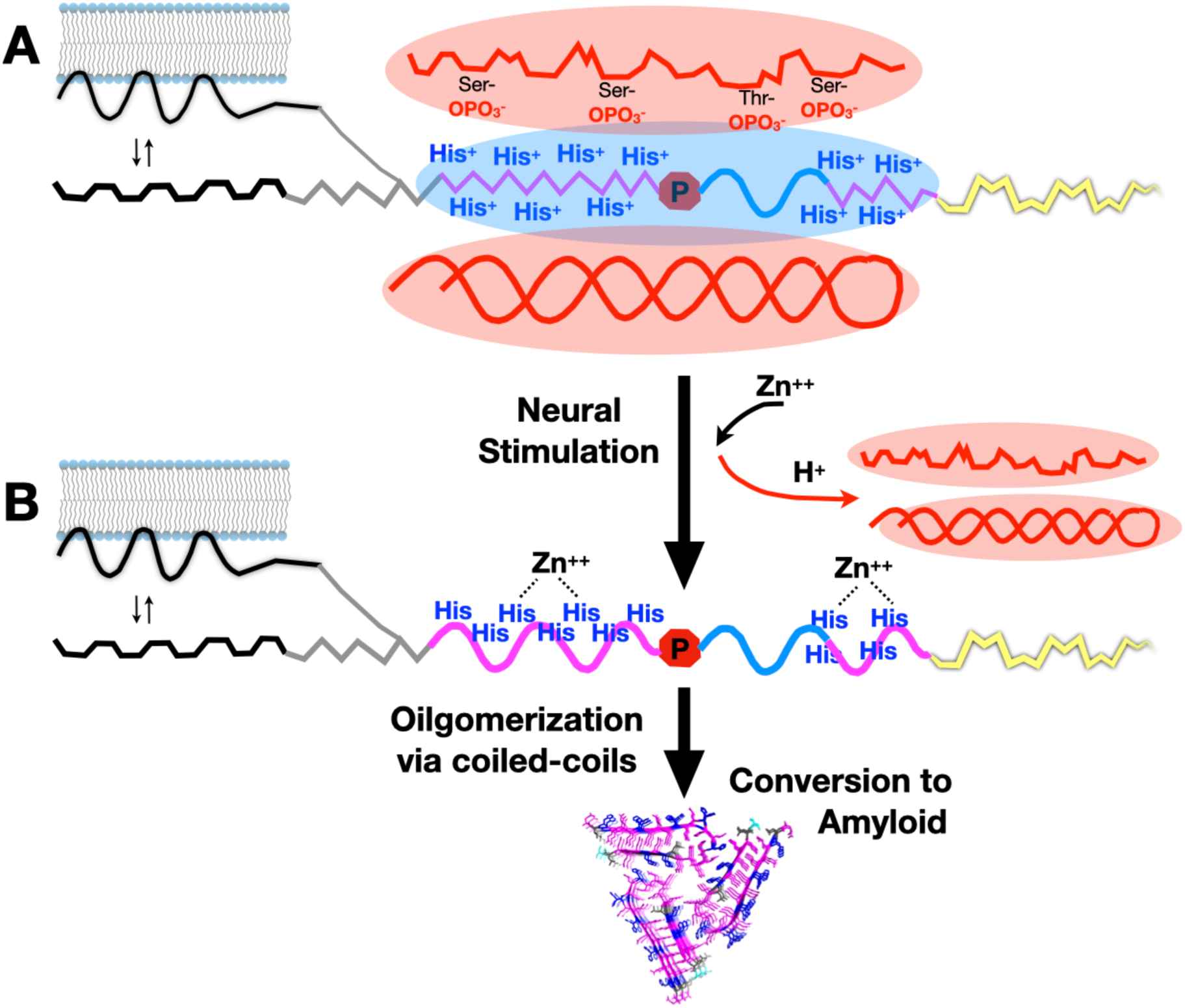
His neutralization unleashes α-helix formation, oligomerization and amyloidogenesis. **A.** The first N-terminal hydrophobic segment (**black**) interacts with membrane which promotes partial α-helix formation as reported by Soria *et al*., (2017). The proximity of phosphorylated Tob protein (top, **red**) RNA (bottom, **red**), and/or possibly other polyanions, creates a negatively charged milieu which favors the cationic form of the eleven His residues (**His +)** of the Q/H-rich segments (**magenta**). This keeps the Q/H-rich segments disordered. Pro 54 (**red** stop sign) would also act to prevent premature amyloidogenesis. The 55-60 residue segment (**blue**) adopts a partial α-helix while the G/S-rich segment (**yellow**) remains disordered and flexible. **B.** Following neural stimulation, the entry of Zn^++^ and other cations could displace H^+^ and bind to the neutral form of His residues (**His**) as proposed by Bajakian *et al*., (2017); releasing Tob and RNA (**red**). The neutral form of His allows partial α-helix formation of the Q/H-rich segments, unlocking pathways to amyloid formation through coiled-coil intermediates.

The role of the initial hydrophobic segment (**Figure 1A**) in Orb2A on amyloidogenesis has been debated. It was found to be important for Orb2 mediated memory, as even the conservative substitution of Phe 5 for Tyr affected activity (Majumdar *et al*., 2012). Moreover, this segment, and not the amyloidogenic Q/H-rich stretch, was found to adopt an α-helix which later evolved into amyloid *in vitro* as monitored by circular dichorism spectroscopy and solid state NMR spectroscopy (Cervantes *et al*., 2016). However, this segment was not observed in the physiological amyloid, which is composed chiefly of Orb2B, formed in *Drosophila* heads (Hervás *et al*., 2020a). Instead, the amyloid is composed of the longer Q/H-rich stretch. Here, this Orb2A N-terminal hydrophobic segment was found to be devoid of detectable populations of secondary structure, at both pH 4.0 and pH 7.0. By constrast, this segment was reported to form an amphiphilic α-helix in the presence of membrane mimetics (Soria *et al*., 2017). Therefore, it is possible that in *in vivo* conditions, this segment may interact with membranes to favor the formation of an α-helix which could promote association events that facilitate functional amyloid formation by the Q/H-rich stretch (**Figure 5**). Once formed, the Orb2A amyloid would nucleate amyloid formation in Orb2B.

The 200-residue long G/S-rich region links the PLD to the folded RRM and ZnF domains of Orb2 (**Sup. Fig. 1A**). Although the physical linkage of the PLD to the RNA-binding domains is required for Orb2 function in memory (Li *et al*., 2016), our characterization of the first 20 residues of this region suggests that it is highly disordered and flexible. Nevertheless, as Orb2 forms granules in cells (Hervás *et al*., 2016) and considering that the disordered region between the PLD and first RRM in human CPEB3 has been recently found to be necessary and sufficient for condensate formation by liquid-liquid phase separation (Ramírez de Mingo *et al*., 2020b), whether or not the G/S-rich region of Orb2 could perform a similar role remains an open question.

### Experimental Procedures

DNA coding for the first 88 residues of Orb2A: M_1_YNKFVNFIC_10_-GGLPNLNLNK_20_-PP<u>QLHQQQHQ_30_-QQHQQHQQHQ_40_-QQQQLHQHQQ_50_-QLS</u>PNLSALH_60_-HHHQQQQQ*LR*_*70*_-*ESGGSHSPSS*_*80*_*-PGGGGGGS*_*88*_, which comprise the PLD domain (underlined) and initial residues of the Gly/Ser-rich region (shown in italics above), was subcloned into a pET28(a) vector containing a TXA fusion protein, a hexaHis tag for IMAC purification and a TEV protease tag. After TEV cleavage, a Gly and Ser residues remained attached to the N-terminus of the Orb2A protein construct. Following expression in minimal media enriched with ^15^NH_4_Cl and ^13^C-glucose as the sole sources of nitrogen and carbon, the resulting protein was purified by Ni^++^ affinity chromatography using a GE Biosciences ÄKTA FPLC system, using the buffer 20 mM TrisHCl, 500 mM NaCl [pH 8.0] adding imidazole (500 mM) for the elution. Next, the protein was cleaved with TEV protease, using 20 mM TrisHCl, 150 mM NaCl, 1 mM DTT, 0.5 mM EDTA, pH 7.0, and re-purified to remove the cut TXA and His tag. Because of the high number of His residues in Orb2A, the protein remained attached to the column and co-eluted with TXA. To remove the fusion protein, we performed anion exchange chromatography using 20 mM TrisHCl, 8 M Urea, 1 mM DTT [pH 7.0], where the elution was achieved applying a linear gradient until 1 M of NaCl was present in the buffer. The resulting protein is electrophoretically pure and was stored in concentrated urea at -80 °C until use. Its purity and identity were subsequently corroborated by NMR spectroscopy.

Just prior to NMR spectroscopy, the urea was removed and the protein was transferred to 1.0 mM deuterated acetic acid buffer containing 15 % D_2_O, pH 4.0 using a PD-10 (GE-Biosciences) desalting column. By refractive index, the sample was found to contain less than 10 mM residual urea. Following NMR spectral acquisition at pH 4.0, the sample was transferred to pH 7.0 by adding a 10X buffer stock of PBS prepared in D_2_O. Upon adding the PBS stock, the initial pH was 6.2; it was increased to 7.01 by adding small alquots of 0.3 M KOH. The final buffer content of the pH 7 sample was 0.9 mM deuterated acetate, 25 mM Na_2_HPO_4_, 25 mM NaH_2_PO_4_, 100 mM KCl and 1 mM TCEP as the reducing agent. The final sample volumes were 0.201 mL (pH 4) and 0.226 mL (pH 7). A final series of 2D and 3D spectra were recorded following the addition of ZnCl_2_ to a final concentration of 2 mM. All spectra were recorded in a water-matched, 5 mm diameter Shigemi NMR tube.

### NMR Spectral Assignments

All NMR spectra were recorded on a Bruker Avance NMR spectrometer operating at 800 MHz (^1^H), except the last series examining the effects of Zn^++^ binding, which were recorded with a Bruker 800 MHz Neo Avance NMR spectrometer. This instrument is equipped with a triple resonance (^1^H, ^13^C, ^15^N) cryoprobe and Z-gradients. The program Topspin (Bruker Biospin) versions 2.1 and 4.0.8 were used to record, process and analyze the spectra. Because the Orb2A’s N-terminal region is disordered and since its sequence is of low complexity and contains many repeated consecutive residues, it presents special challenges for NMR spectral assignment. Therefore, instead of the standard approach based by ^1^HN excitation and ^13^Cα and ^13^Cβ connectivities (Sattler *et al*., 1999), we have applied an unconventional proton-less approach based on a pair of 3D ^13^C-detected spectra which provide consecutive ^13^CO and ^15^N backbone connectivities as these nuclei retain more dispersion in IDPs (Pantoja-Uceda & Santoro, 2014). Even with this approach, some cases of sequential residues with identical ^13^CO and ^15^N chemical shift values were observed. Therefore, to overcome these ambiguities, an additional 3D ^1^H-detected spectrum which yields consecutive ^1^HN-^1^HN connectivities was recorded (Sun *et al*., 2005; Pantoja-Uceda & Santoro, 2013). The analysis of these data led to the essentially complete spectral assignment at pH 4.0 as described in the **Results** section. After measuring relaxation measurements at pH 4.0, the sample pH was increased to pH 7.0. As described in the **Results** section, the sample tends to aggregate at neutral pH. Because of its lower solution concentration, the assignment was carried out by conventional ^1^H-detected 2D ^1^H-^15^N HSQC and 3D CBCAcoNH, HNCO and HNCA spectra and comparison with the pH 4 spectra. A list of the NMR experiments and their parameters is shown in **Sup. Table 1**. ^1^H peak widths in ^1^H-^15^N HSQC spectra were measured using a Gaussian function in the program NMRFAM Sparky 1.4. ^3^J_HNHA_ coupling constants were calculated based on the relative signal intensities of the ^1^HN/^1^Hα crosspeak : ^1^HN/^1^HN diagonal peak using either peak integration in Topspin 4.0 or peak heights in NMRFAM Sparky 1.4 as previously described (Vuister & Bax, 1993).

The expected chemical shift values for the ^13^Cα, ^13^Cβ, ^1^Hα, ^1^HN, ^15^N and ^13^CO nuclei in statistical coil ensembles were calculated using the parameters tabulated by Kjaergaard and Poulsen (2011) and Kjaergaard *et al*., (2011), and implemented on the https://spin.niddk.nih.gov/bax/nmrserver/Poulsen_rc_CS/ server at the Bax laboratory. These values were subtracted from the experimentally measured chemical shift values (δ) to calculate conformational chemical shifts (Δδ).

### NMR Relaxation Measurements

The dynamics on the ps–ns time scales were probed by measuring the heteronuclear ^15^N{^1^H} NOE (hNOE) of backbone amide groups as the ratio of spectra recorded with and without saturation in an interleaved mode. An eleven second recycling delay was employed. Uncertainties in peak integrals were estimated from the standard deviation of intensities from spectral regions lacking signal and containing only noise. R_1_ρ relaxation rates, which sense dynamic processes on slower μs-ms timescales, were measured by recording two sets of ten ^1^H-^15^N correlation spectra with relaxation delays at 8, 300, 36, 76, 900, 100, 500, 156, 200 and 700 ms. The R_1_ρ relaxation rates were calculated by least-squares fitting of an exponential decay function to peak integral data, which were obtained using Topspin 4.0.8.

### Bioinformatic Analysis

Disorder predictions for the Orb2A sequence were performed with PONDR-XSL2 and PONDR-XL1-XT (Xue *et al*., 2010).

## Abbreviations Used

CPEB: Cytoplasmic Polyadenylation Element Binding protein.
Orb2A: The *Drosophila* protein encoded by the <u>o</u>o18 <u>R</u>NA-<u>b</u>inding (*orb*) gene, predominantly neuronal isoform 2A.
IDR: Intrinsically Disordered Region (Residues 1-292 of Orb2A)
NMR: Nuclear Magnetic Resonance
PLD: Prion-Like Domain (Residues 1-68 of Orb2A)

## Acknowledgements

JO was supported by a Leonardo Grant (BBM_TRA_0203) from the BBVA Foundation. The authors acknowledge FCT-Portugal for the PhD studentship attributed to Sara S. Félix (PD/BD/148028/2019). This study was supported by project SAF2016-76678-C2-2-R (DVL) from the Spanish Ministry of Economy and Competitivity. NMR experiments were performed in the “Manuel Rico” NMR Laboratory (LMR) of the Spanish National Research Council (CSIC), a node of the Spanish Large-Scale National Facility (ICTS R-LRB). This work was also partially supported by the Applied Molecular Biosciences Unit – UCIBIO, which is financed by national funds from the Portuguese FCT (UIDB/04378/2020). We are grateful to Daniel Ramírez de Mingo, Rubén Hervás and Mariano Carrión-Vázquez for critical comments on the manuscript, and to José Manuel Peréz Cañadillas for the pET28-TXA-His cDNA vector.

---

**Sup. Table 1:** NMR Spectral Parameters.

| Experiment | Number of Scans | Sweep Width (ppm) | Matrix |
| --- | --- | --- | --- |
| *Assignment pH 4.0* |  |  |  |
| 1D $^1\text{H}$ | 8 | 10 | 32k |
| 2D $^1\text{H}$-$^{15}\text{N}$ HSQC | 4 | 10 $^1\text{H}$ x 20 $^{15}\text{N}$ | 2k x 512 |
| 2D CON* | 4 | 11 $^{13}\text{CO}$ x 35 $^{15}\text{N}$ | 1k x 512 |
| 3D HNCO | 8 | 10 $^1\text{H}$ x 20 $^{15}\text{N}$ x 11 $^{13}\text{C}$ | 2k x 64 x 128 |
| 3D hacacoNcaNCO* | 8 | 11 $^{13}\text{CO}$ x 35 $^{15}\text{N}$ x 20 $^{15}\text{N}$ | 1k x 48 x 96 |
| 3D hacaCONcaNCO* | 8 | 11 $^{13}\text{CO}$ x 35 $^{15}\text{N}$ x 21 $^{13}\text{CO}$ | 1k x 48 x 96 |
| 3D HncocaNH | 8 | 10 $^1\text{H}$ x 20 $^{15}\text{N}$ x 10 $^1\text{H}$ | 2k x 96 x 48 |
| 3D CCC...CON* | | 11 $^{13}\text{CO}$ x 35 $^{15}\text{N}$ x 60<br>$^{13}\text{C}(\text{aliphatic})$ | 1k x 56 x 96 |
| 3D CBCACON* | 8 | 11 $^{13}\text{CO}$ x 35 $^{15}\text{N}$ x 70<br>$^{13}\text{C}(\text{aliphatic})$ | 1k x 64 x 80 |
| 3D HNCA | 4 | 11 $^1\text{H}$ x 20 $^{15}\text{N}$ x 25 $^{13}\text{C}$ | 2k x 48 x 128 |
| 3D HNHA<sup>§</sup> | 8 | 11 $^1\text{HN}$ x 23 $^{15}\text{N}$ x 11 $^1\text{H}\alpha$ | 2k x 96 x 48 |
| *Assignment pH 7.0 (both before and after adding $\text{ZnCl}_2$)* |  |  |  |
| 2D $^1\text{H}$-$^{15}\text{N}$ HSQC | 4 | 10 $^1\text{H}$ x 20 $^{15}\text{N}$ | 2k x 512 |
| 3D HNCO | 16 | 10 $^1\text{H}$ x 20 $^{15}\text{N}$ x 11 $^{13}\text{C}$ | 2k x 32 x 64 |
| 3D HNCA | 12 | 10 $^1\text{H}$ x 20 $^{15}\text{N}$ x 20 $^{15}\text{N}$ | 2k x 64 x 90 |
| 3D CBCAcoNH | 16 | 10 $^1\text{H}$ x 20 $^{15}\text{N}$ x 70 $^{13}\text{C}$ | 2k x 32 x 64 |
| *Relaxation* |  |  |  |
| 2D $\{^1\text{H}\}$-$^{15}\text{N}$ NOE<sup>¶</sup> | 8-16 | 10 $^1\text{H}$ x 20 $^{15}\text{N}$ | 2k x 512 |
| 2D $^1\text{H}$-$^{15}\text{N}$ T<sub>1</sub>ρ<sup>‡</sup> | 8-16 | 10 $^1\text{H}$ x 20 $^{15}\text{N}$ | 2k x 256 |

\* $^{13}\text{C}$ detection; processed with in-phase anti-phase (IPAP) virtual decoupling.
§ Recorded to measure $^1\text{HN}$ - $^1\text{H}\alpha$ coupling constants.
¶ A delay of 11 seconds between pulses was used. Recorded in interleaved mode.
‡ Ten spectra with delays of 8, 300, 36, 76, 900, 100, 500, 156, 200 and 700 ms were recorded.

**Sup. Fig. 1:**
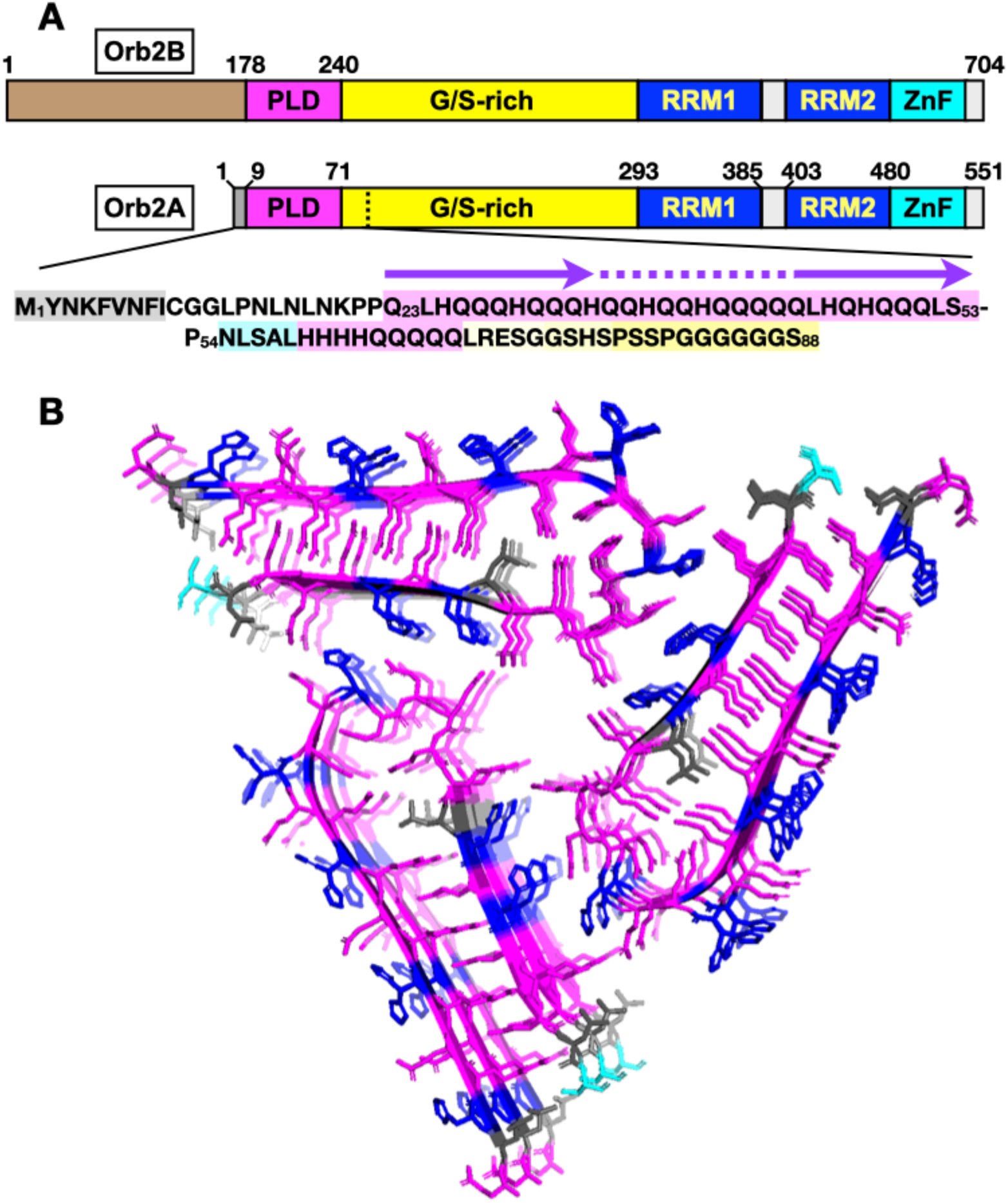
Orb2 domain and amyloid core structure. **A.** Orb2A and Orb2B differ at the N-terminus (**brown** versus **gray** regions) but both contain a PLD (**magenta**) two RRMs, and a ZZ-type ZnF domain (**torquoise**). The sequence of the OrbA region studied here includes a hydrophobic N-terminal segment (shaded **gray**), a Q/H-rich amyloid-forming segment (shaded light **magenta**), a short segment prone to form α-helix (shaded **blue**), a second modest H·Q-rich segment (shaded light **magenta**), and the beginning of the G/S-rich region (shaded **yellow**). The purple arrows and dotted lines mark residues adopting β-strands and a turn, respectively, in the amyloid core structure shown in panel **B**. **B.** Structure of the Orb2 amyloid core (**PDB: 6VPS**), solved by cryo-EM (Hervás *et al*., 2020a). Residue color core: Q = **magenta**, H = **blue**, L = **gray**, S = **cyan**.

**Sup. Figure 2.**
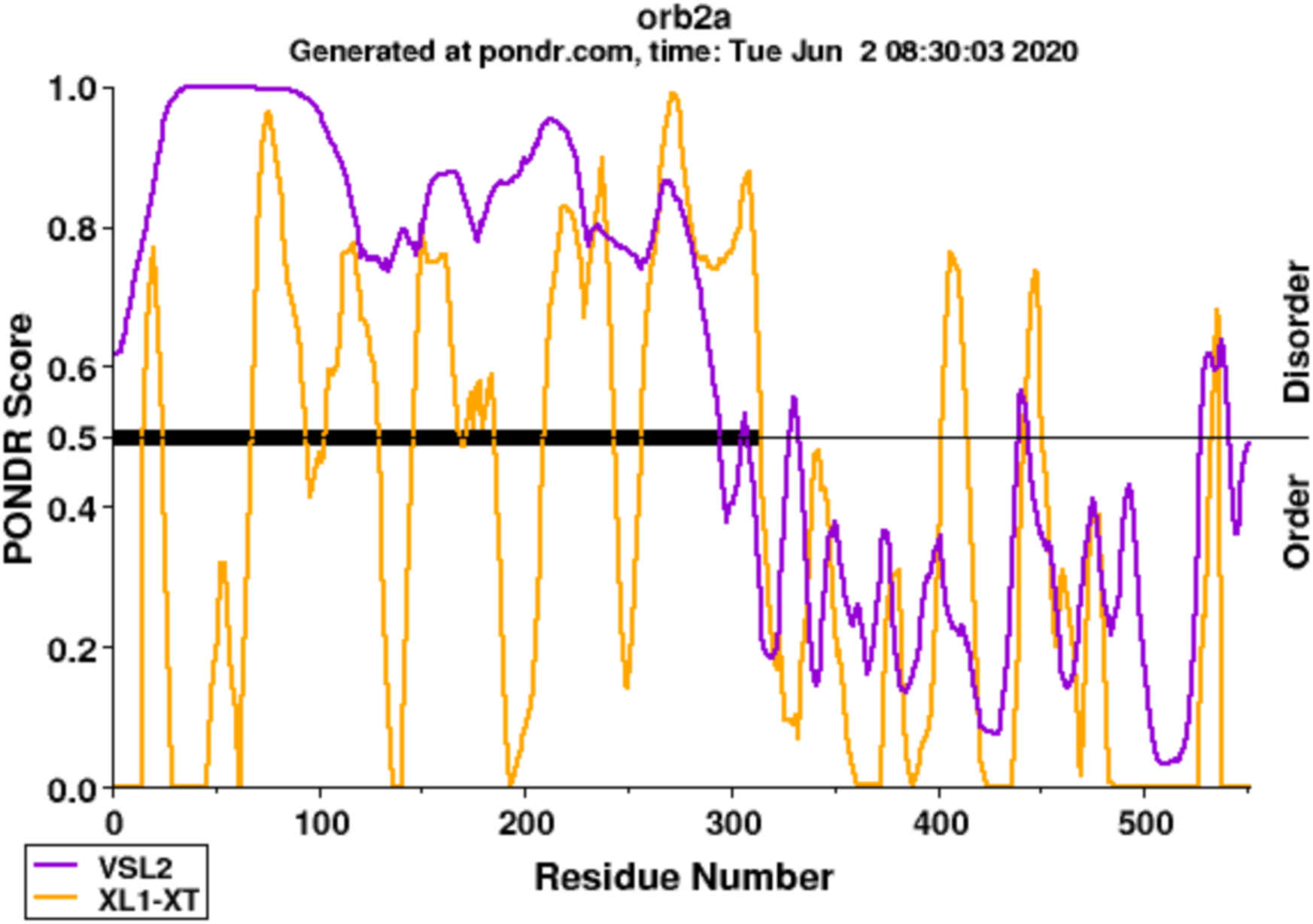
Predicted Regions of Order and Disorder in Orb2A (Related to Sup. Fig 1A and the Introduction in the main text) Analysis with the program PONDR predicts that Orb2A’s PLD and G/S-rich regions will be unfolded. The folded RRM and ZnF domains, which begin near residue 300 (as shown in **Sup. Fig. 1A**) are correctly predicted to be ordered.

**Sup. Figure 3.**
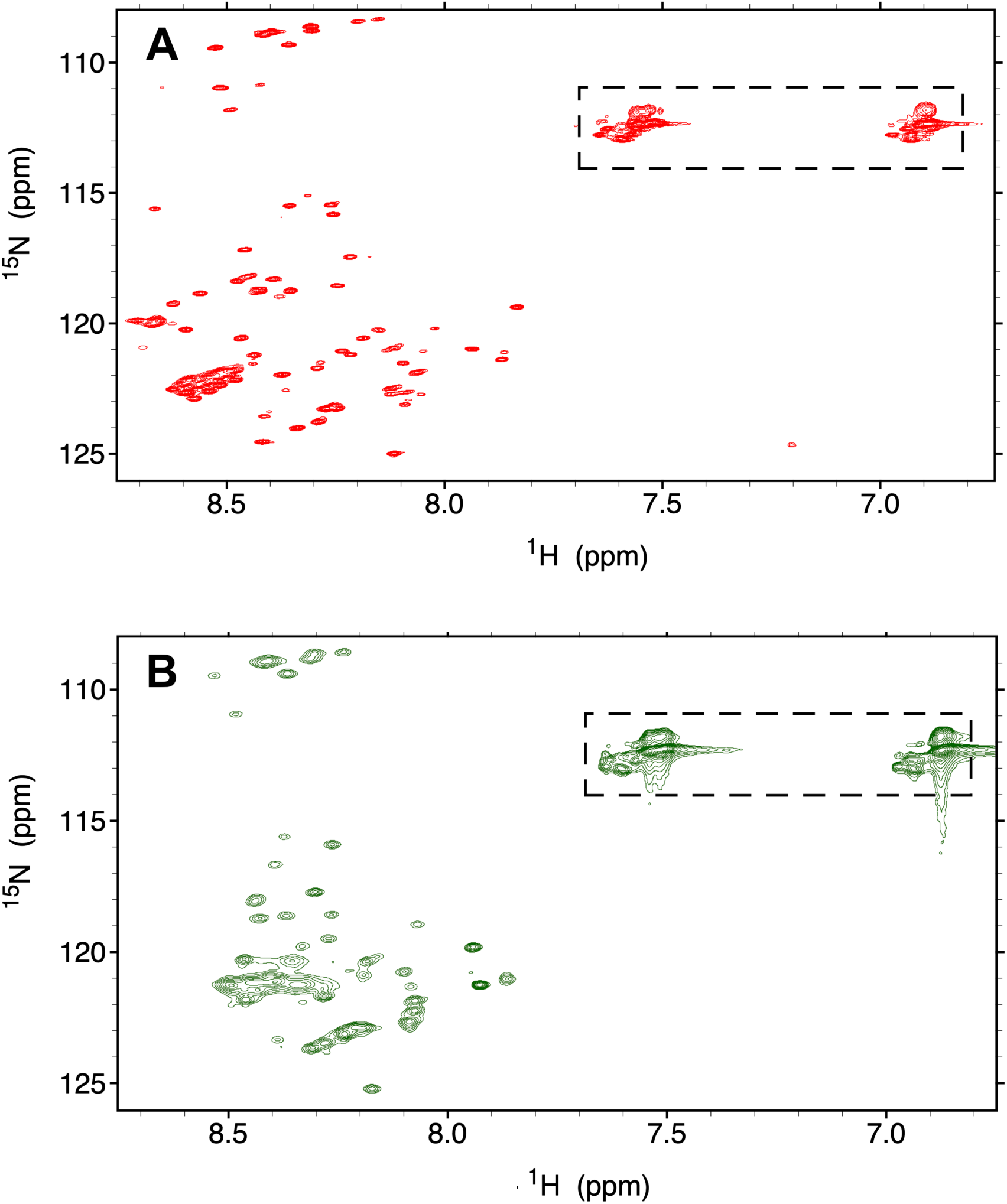
(Related to Figure 1A and 3A in the Main Text) Full 2D ^1^H-^15^N HSQC spectra of Orb2A PLD with the Asn/Gln side chain ^1^H_2_-^15^N region. **A.** 25°C, pH 4, 1 mM deuterated acetic acid. The Asn/Gln side chain H_2_N-signals are boxed. **B.** 25°C, pH 7, in PBS buffer. The Asn/Gln side chain H_2_N-signals are boxed.

**Sup. Figure 4.**
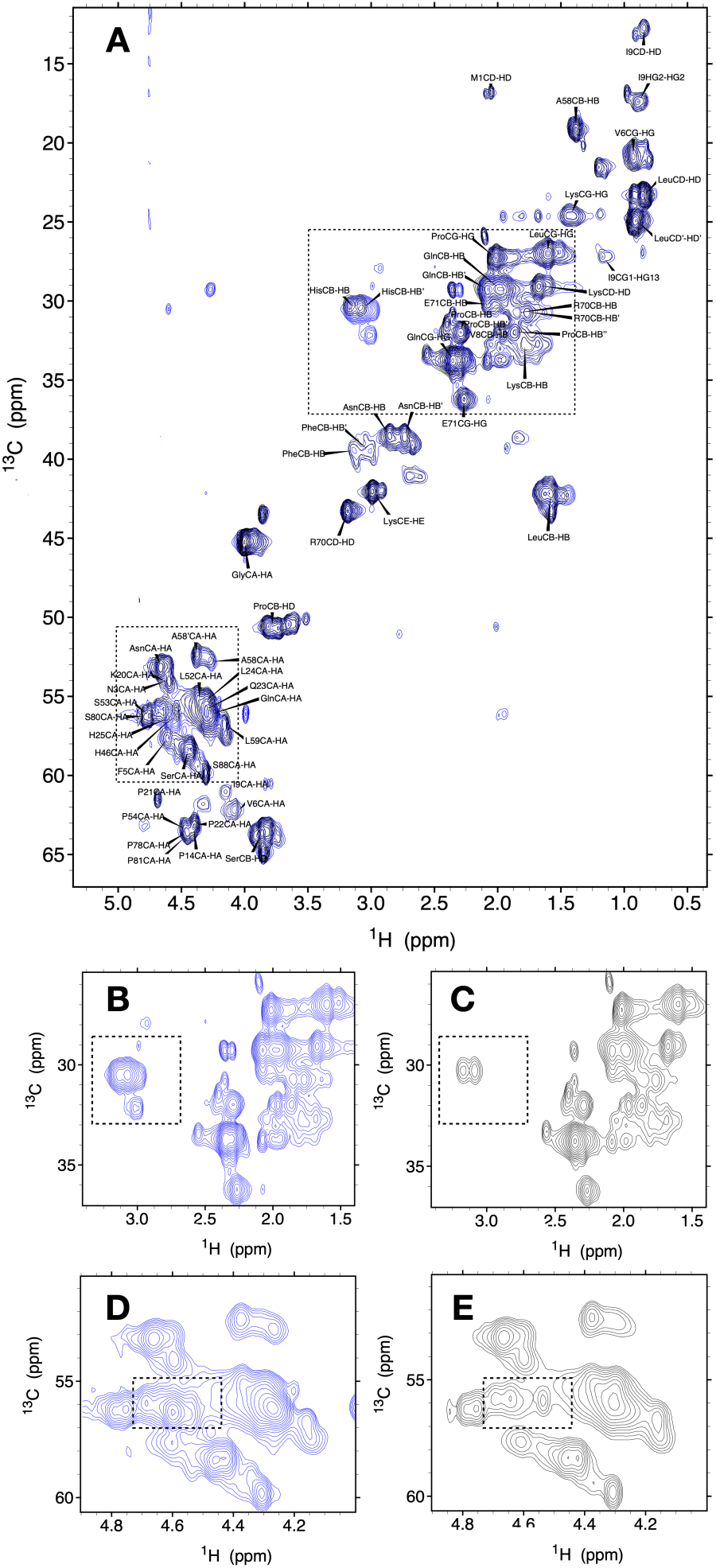
^1^H-^13^C HSQC spectral evince that His residues of Orb2 PLD bind zinc chloride. (Related to Figures 4 and 5 in the Main Text) **A**. Complete, assigned ^1^H-^13^C HSQC spectrum at 25°C, pH 7 in PBS without zinc chloride (**blue**) and with zinc chloride (**black**). Zoomed views of the boxed areas are shown in panels **B, C, D** & **E**. **B** & **C**. Zoomed view of panel **A**; the boxed area here highlights the histidine ^1^Hβ-^13^Cβ correlations before (**B**) and after (**C**) ZnCl_2_ addition. **D** & **E**. Zoomed view of panel **A**; the boxed zone shows the histidine ^1^Hα-^13^Cα correlations prior to (**D**) and following (**E**) ZnCl_2_ addition.

## Notes

http://www.bmrb.wisc.edu

### Competing Interest Statement

The authors have declared no competing interest.

### Summary of Updates

The revised manuscript includes some additional grant information. Everything else is the same.

